# QuickProt: A Fast and Accurate Homology-Based Protein Annotation Tool for Non-Model Organism Genomes

**DOI:** 10.1101/2025.03.28.645852

**Authors:** Guisen Chen, Hehe Du, Zhenjie Cao, Ying Wu, Chen Zhang, Yongcan Zhou, Jingqun Ao, Yun Sun, Zihao Yuan

## Abstract

Despite the increasing number of genomes, the annotation of protein-coding genes is still lacking. In some genomes where transcriptome data is not available, various tools have been developed to annotate genes based on homologous proteins, but each tool has its limitations. Here, we propose quickprot, a user-friendly command-line tool that constructs a set of redundancy-free gene models from the target genome based on the homologous protein sequences of a set of closely related species. We tested its applicability in stonefish, and the results showed that it has high accuracy and runs faster than existing tools. This low-cost, high-precision annotation method is expected to become a useful tool for genome annotation, which will contribute to the research of comparative genomics. The software is implemented in Python and licensed under the MIT License. The source code, documentation, and tutorials can be obtained at https://github.com/thecgs/quickprot.

## Introduction

With the continuous advancement of genome sequencing technologies, an increasing number of species genomes have been sequenced and published. However, based on the statistical information from NCBI Genomes (https://www.ncbi.nlm.nih.gov/datasets/genome/) as of March 12, 2025, only approximately 18% of eukaryotic genomes have completed gene set annotation. This gap has significantly hindered the advancement of comparative genomics and related studies on gene structure and functional annotation. Genome annotation remains a bottleneck due to the lack of efficient and universally applicable methodologies. As highlighted by the developers of GALBA (Brůna, et al., 2023), one of the major issues in gene annotation is caused by the limitations of computational and human resource faced by major genome repositories (e.g., EBI, NCBI, and Ensembl), which prevent them from applying internal annotation pipelines to all submitted genomes. Furthermore, existing annotation pipelines exhibit inconsistency in their performance across various species, making their universal use challenging for smaller and inexperienced research groups. Additionally, RNA-seq data accessibility is yet another critical challenge, primarily limited by sample or funding acquisitions.

In the lack of transcriptomic evidence, annotating novel genomes by leveraging splice-alignment information from proteins of related species to the target genome constitutes a standard procedure in genomic annotation. Various tools have been developed to perform the protein annotation by aligning proteins to whole genomes, such as blastp (Camacho, et al., 2009), Spaln2 (Iwata and Gotoh, 2012), GenomeThreader (Gremme, 2005), and GeMoMa (Keilwagen, et al., 2019). Nevertheless, those software are CPU intensive and can only align hundreds of proteins per CPU hour. This has caused the alignment of a medium-sized genome to take up to days, making the protein to genome pipelines label and time-consuming.

As a newer alternative, Miniprot (Li, 2023) is a recently developed rapid protein-to-genome aligner that achieves accuracy comparable to established tools. However, miniprot aligns each protein sequence separately, lacking the capacity to consolidate identical gene models or resolve conflicts when multiple proteins are mapped to the same genomic locus. Similar limitations are found in genewise (Birney, et al., 2004), genblastG (She, et al., 2011), and Exonerate (Slater and Birney, 2005), which do not systematically integrate overlapping alignments or address inconsistencies due to redundant genomic annotations. Integrated annotation pipelines such as BRAKER3 (Gabriel, et al., 2024) and MAKER2 (Holt and Yandell, 2011), which combine *ab initio* prediction, homology-based protein alignment, and transcriptome-supported evidence integration, may exhibit limited applicability for genome-wide annotation in scenarios lacking RNA-seq data. The development of BRAKER2 (Brůna, et al., 2021) and GALBA tried to solve the aforementioned challenges by incorporating self-training algorithms, such as GeneMark-ES (Lomsadze, et al., 2005) or AUGUSTUS (Stanke, et al., 2006). Those methods enhance gene structure prediction in protein evidence-depleted genomic regions. However, these approaches present two significant challenges: (1) computational cost escalation due to iterative model refinement and (2) overfitting risks leading to reduced generalizability in taxon-specific contexts. For instance, in the longtooth grouper (*Epinephelus bruneus*) genome analyzed in this study, the predicted gene set exhibited elevated false-positive rates and excessive fragmentation. Such fragmented genes pose critical challenges in comparative genomics, as it may potentially misrepresent gene family expansion patterns or generate spurious hypotheses about taxonomically unique genes due to artifactual open reading frames.

To overcome these challenges, here, we present quickprot (https://github.com/thecgs/quickprot), a new homology-based protein annotation pipeline designed for rapid construction of non-redundant gene models in genome annotation. Taking protein sequences from one or multiple reference species and the target genome as input, our method achieves superior computational efficiency while maintaining high accuracy compared to BRAKER2 and GALBA, as demonstrated by comprehensive benchmarking analyses. The fully open-source nature of this pipeline enables efficient generation of artifact-free gene models, thereby advancing research in comparative genomics, gene structure evolution, and functional annotation in non-model organisms.

## Algorithm principle

The implementation of the quickprot algorithm is mainly divided into three parts (Fig.1A). Firstly, it uses miniprot (v0.12) to align homologous protein sequences with the target genome and expands these high-quality protein coding regions, integrating and assembling them into pseudo-transcripts (lacking the UTR part). Then, it uses TransDecoder (v5.7.1) (Haas, 2024) to predict the coding regions within these pseudo-transcripts. Subsequently, we filtered out low-quality gene models. For the very few that were affected by overlapping genes, resulting in a pseudo-transcripts being composed of multiple transcripts, we based our ORF prediction scores, the size of the overlapping region between two transcripts, and the length of the coding region of the transcript, to carry out effective splitting, ultimately generating a high-precision, non-redundant gene set. In simple terms, the core principle of quickprot is similar to the blotting method. It predicts gene models by marking the coding regions in the target genome sequences with homologous proteins from closely related species. The advantage of this approach is that it reduces computational cost without introducing false positive genes. However, its disadvantage is also obvious. It relies on more comprehensive and accurate homologous proteins to try to find all potential coding regions. Unlike BRAKER2 and GALBA, it cannot predict coding regions outside homologous proteins. But with the further improvement of protein databases such as SwissProt and OrthoDB, we believe the impact of this drawback will be further reduced.

**Fig. 1.**
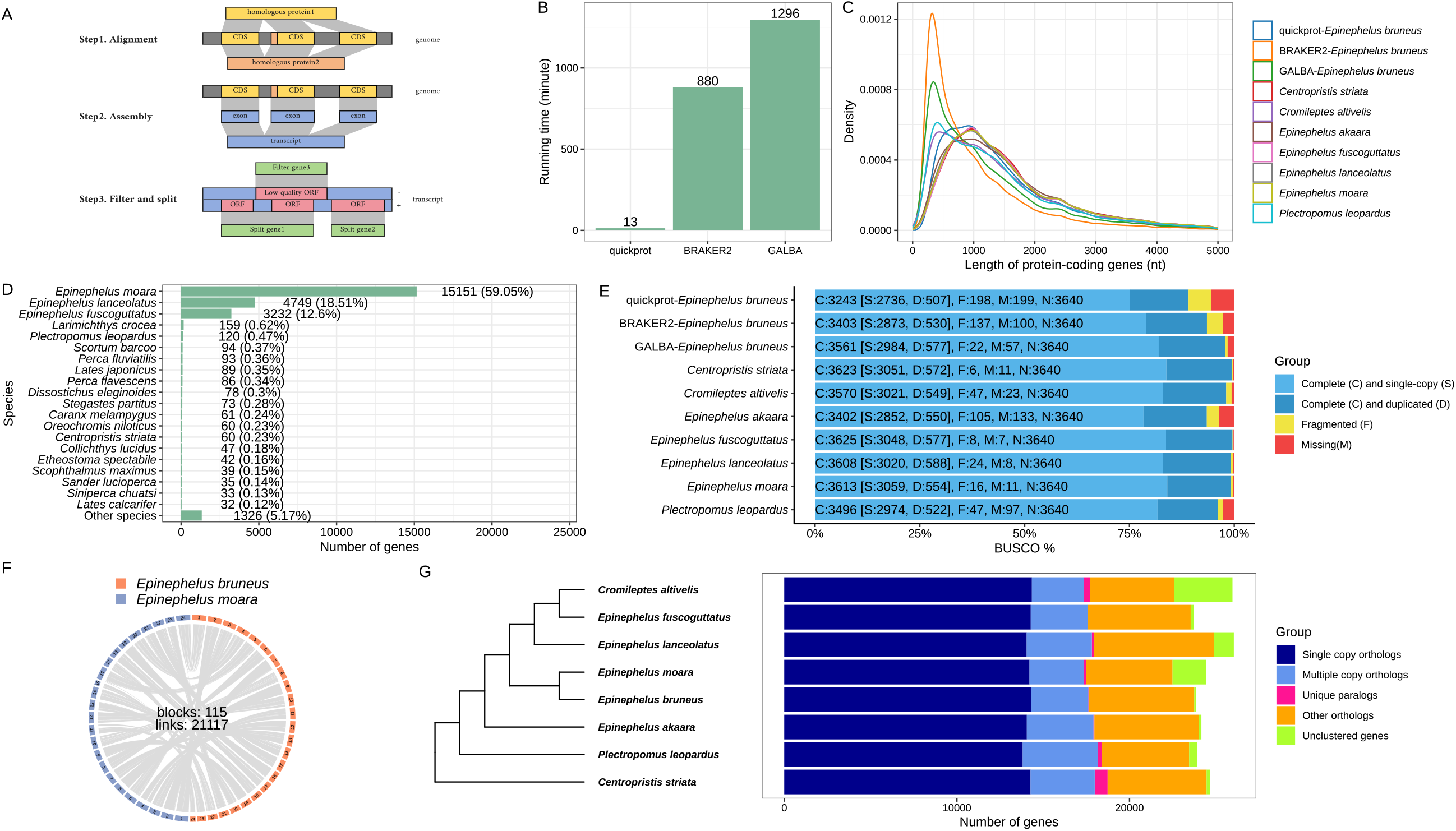
Algorithm principle and technical verification of quickprot. A: Schema of quickprot algorithm. B: In Ubuntu 22.04, the system uses 24 threads of quickprot, BRAKER2, and GALBA process runtime. C: Density distribution of coding gene length. D: NR-annotated species distribution using quickprot-predicted longtooth grouper protein-coding genes. E: BUSCO evaluation. F: Genome synteny between longtooth grouper and kelp grouper. G: Orthologous genes and species tree between longtooth grouper and seven other teleost species.

## Technical validation

To evaluate the reliability of our method, we conducted technical validation using the genome of the longtooth grouper, a commercially significant marine fish species in East Asia belonging to the subfamily Epinephelinae. While a genome assembly for this species is publicly available (GCA_041684075.1), no annotated gene models and transcriptomic datasets exist. In this study, we downloaded the protein sequences of seven closely related species from NCBI, Dryad, Figshare, and NGDC, including the giant grouper (*E*.*lanceolatus*, GCF_005281545.1), tiger grouper (*E*. *fuscoguttatus*, GCF_011397635.1), kelp grouper (*E. moara*, GCF_006386435.1), black sea bass (*Centropristis striata*, GCF_030273125.1), leopard coral grouper (*Plectropomus leopardus*, https://ngdc.cncb.ac.cn/gwh/Assembly/29542/show), and humpback grouper (*Cromileptes altivelis*, https://figshare.com/articles/dataset/humpback_grouper_genome/24145230/2), and hong kong grouper (*E*.*akaara*, doi:10.5061/dryad.4398b9f). As well as the genome sequences of the longtooth grouper and the kelp grouper. The gene sets of all species were filtered using the get_longest_transcript_gff3.py script (integrated in quickprot) to remove redundant transcripts and retain the longest transcripts. Then, under the Ubuntu-22.04 system, we tested the BRAKER2, GALBA, and quickprot pipeline using 24 threads and obtained 39,381, 55,416, and 26,046 protein-coding genes, respectively. According to the statistics of the gene sets of seven closely related species, the actual number of genes should be in the range of 23,722-25,965. The results show that BRAKER2 and GALBA will introduce more false positives, while quickprot seems to perform better in controlling false positives. Moreover, quickprot runs very fast. In the study, quickprot is about 67.69 times faster than BRAKER2 and about 99.69 times faster than GALBA (Fig.1B). To explore whether the prediction results of BRAKER2 and GALBA are more fragmented, we counted the density distribution of the length of protein-coding genes predicted by BRAKER2, GALBA, and quickprot, as well as those of seven other closely related teleost fish, as shown in Fig.1C. The protein-coding genes predicted by BRAKER2 and GALBA showed a significant peak around 325 nt, indicating that the gene lengths predicted by BRAKER2 and GALBA are mostly concentrated around 325 nt. On the other hand, the results predicted by quickprot are more similar to the length distribution patterns of the other seven teleost fish, showing more accurate prediction performance than BRAKER2 and GALBA.

To rigorously assess the accuracy of quickprot predictions, we performed sequence similarity searches against the Non-Redundant (NR) protein database using DIAMOND (v2.1.8.162) (Buchfink, et al., 2021) under strict alignment criteria (E-value threshold = 1e-10). The results demonstrated that 25,627 out of 26,046 predicted protein-coding genes (98.39%) showed significant matches to known protein sequences, indicating high annotation reliability. A statistical analysis of the species distribution in the NR database shows that (Fig.1D) the species most closely related to the longtooth grouper is the kelp grouper. In the early stages, the morphological characteristics and distribution of these two species were very similar, so there were many controversies about their classification. Some researchers regarded them as one species, while most others divided them into two species based on morphological characteristics and PCR amplification of the ND2 gene and ITS1 region, as well as cytogenetic analysis (Guo, et al., 2009; Guo, et al., 2014). For this, we further conducted a synteny analysis of the two species, which was completed using JCVI (v1.4.23) (Tang, et al., 2024), with a c-score value set to 0.99 and a minspan of 30. We also used OrthoFinder (v2.5.5) (Emms and Kelly, 2019) to identify orthologous genes and inferred a rooted species tree using the MSA (multiple sequence alignment) method. The NR annotation results show that there is a 59.05% high identity between the protein-coding genes of the two species (Fig. 1D), and the synteny results show that the two species have identified a total of 21,117 homologous gene pairs, and 115 genomic collinear regions (Fig. 1F). This reflects that the protein-coding genes of the two species have both a high similarity and differences. OrthoFinder analysis revealed that these closely related species have highly similar structures and compositions of orthologous and paralogous genes (Fig. 1G). The phylogenetic tree of species suggests that longtooth grouper and kelp grouper are the most closely related species, and the topology of the species tree is highly consistent with some early species trees constructed from nuclear and mitochondrial genomes (Cao, et al., 2022; Zhuang, et al., 2013). These results indirectly support the hypothesis that longtooth grouper and kelp grouper are two species, and at the same time verify the reliability of the prediction results of quickprot.

To further validate the predictive performance, we conducted BUSCO (Benchmarking universal single-copy orthologs) assessments using compleasm (v0.2.6) (Huang and Li, 2023) on protein-coding genes predicted by BRAKER2, GALBA, and quickprot, along with those from seven phylogenetically proximate species. The analysis was performed under the actinopterygii_odb10 lineage dataset. The BUSCO results show (Fig.1E) that GALBA performed the best in this test (97.83%), followed by BRAKER2 (93.49%), while quickprot only achieved an 89.08% completeness. This is partly because the quickprot design lacks *de novo* prediction modules, limiting its ability to identify coding regions beyond conserved orthologs. On the other hand, the miniprot stringent alignment criteria (>95% sequence identity) prioritize precision at the cost of sensitivity. Users may adjust these thresholds based on the evolutionary divergence between reference proteins and the target species. Although BUSCO is widely adopted as a standard for proteome completeness assessment, the superior BUSCO scores of BRAKER2 and GALBA come at the expense of increased false positives. As noted by the BRAKER3 developers (Gabriel, et al., 2024), the design of BUSCO does not penalize false-positive gene predictions nor detect structural errors (e.g., split/fused genes or incorrect exon boundaries). Consequently, BUSCO scores should be interpreted as rough indicators of prediction sensitivity rather than comprehensive measures of gene structure accuracy or completeness.

## Discussion

In this study, we proposed quickprot, a new tool for predicting genome protein-coding genes based on homologous proteins. In the non-model animal longtooth grouper, which applied in this study, we showed that quickprot had better performance compared with BRAKER2 and GALBA in terms of computational efficiency and false-positive control. However, despite those strengths, compared to the current reference annotation process, quickprot still has its limitations. After assembling transcripts with quickprot, TransDecoder is used to identify candidate coding regions. Quickprot, therefore, inherits some shortcomings of TransDecoder, specifically when a small number of homologous proteins are given as input. This may potentially lead to very few assembled transcripts since TransDecoder needs hundreds of candidate sequences in order to train species-specific models (Haas, 2024). Moreover, the accuracy of quickprot relies on more comprehensive homologous proteins that will cover every potential coding region. In contrast to BRAKER2 and GALBA, which incorporate *de novo* gene prediction components, quickprot won’t be capable of detecting genes with low homology to the input proteins. Although the addition of *de novo* gene prediction can help to boost the gene detection sensitivity, it also has a greater risk of introducing false positives. (Brůna, et al., 2021; Brůna, et al., 2023; Gabriel, et al., 2024). Furthermore, another drawback of adding *de novo* prediction training is that it requires longer running time and higher computational resources. As with all those annotation tools, there is no perfect solution or tool for all species, and the best strategy is dictated by the research needs and available resources. Luckily, as sequencing technologies improved, increasingly representative species of major clades have been sequenced for their genomes and proteomes released to the public. Which will provide more options for choosing the closest homologous species as input. In this case, Quickprot shows great advantages in the prediction speed of protein annotation and also takes into account high accuracy, although its sensitivity is slightly inferior to BRAKER2 and GALBA. However, it is undeniable that this low-cost, high-accuracy annotation method can be expected to be a useful tool for genome annotation, facilitating the research of comparative genomics, such as incorporating more species to a wider phylogenetic analysis at a lower cost.yw

## Acknowledgements

Financial supports for this study were provided by the National Natural Science Foundation of China (U22A20534) and the Innovational Fund for Scientific and Technological Personnel of Hainan Province (KJRC2023C38).

## Author contributions

Conceptualization: Y.S and Z.Y.; Software, Formal analysis, Validation, and Visualization: G.C; Supervision: H.D., Z.C., Y.W., C.Z., Y.Z., J.A; Resources: Y.Z.; Funding acquisition: Y.S. All authors contributed to the article and approved the submitted version. All authors have read and agreed to the published version of the manuscript.

